# Effects of vertical sleeve gastrectomy prior to pregnancy on bone mass, microarchitecture and material properties in the female rat

**DOI:** 10.1101/2024.05.22.595335

**Authors:** Malory Couchot, Françoise Schmitt, Morgane Mermet, Céline Fassot, Guillaume Mabilleau

**Affiliations:** Univ Angers, Nantes Université, ONIRIS, Inserm, RMeS, UMR 1229, F-49000, Angers, France; Univ Angers, HIFIH, F-49000 Angers, France; CHU Angers, Paediatric department, 49933 Angers, France; Univ Angers, Inserm, CNRS, MITOVASC, F-49000 Angers, France; CHU Angers, Cell and tissue pathology, 49933 Angers, France

**Keywords:** vertical sleeve gastrectomy, bone mass, bone microarchitecture, bone material properties, pregnancy, lactation

## Abstract

Obesity is a major public health issue worldwide. Despite various approaches to weight loss, the most effective technique for reducing obesity, as well as diabetes and associated diseases, is bariatric surgery. Increasingly, young women without children are undergoing bariatric surgery, vertical sleeve gastrectomy (VSG) being the most common procedure nowadays. However, despite several reports suggesting bone loss after VSG, little is known about the potential additive effects of gestation and lactation after VSG to bone health. This study investigated the combined effects of pre-gestational VSG and subsequent gestation/lactation on bone metabolism in a rat model fed a high fat high sugar (HFHS) diet, with a focus on bone biomechanics, mass, microarchitecture and material properties. Furthermore, bone mass and remodelling was followed longitudinally by microCT prior to surgery, 4 weeks post-surgery, after weaning and at sacrifice. Significant alterations in bone mass and microarchitecture, characterized by changes in trabecular thickness and number, as well as changes in bone formation and resorption were influenced by both surgery and reproductive demands. Mechanical testing at sacrifice demonstrated compromised long bone fragility, in rat with HFHS regardless of the surgical procedure (Sham or VSG). Furthermore, analysis of bone material properties highlighted potential disruptions in the pattern of bone mineralization in sham and VSG animals fed a HFHS diet. These findings underscore the complex interplay between pre-gestational VSG and subsequent gestation/lactation in modulating bone metabolism. Understanding these combined effects is essential for optimizing surgical strategies and developing targeted interventions to mitigate potential bone-related complications associated with VSG in reproductive-aged individuals.

## 1. INTRODUCTION

Obesity is a growing epidemic worldwide, with numerous health complications and associated problems including bone frailty (1). Obesity is defined as a body mass index (BMI) ≥ 30 and is thought to affect nowadays around 890 million adult individuals and 160 million children worldwide (2). A bone fragility is thought to occur in children, but it is less clear whether this also happens in adult individuals(3). Although excessive weight can augment bone mass and mineral density, it also increases the risk of fracture at specific sites (4).

The current management of obese individuals is represented by lifestyle changes and therapeutic monitoring, accompanied by an increase in physical activity (5). Pharmacological treatment may also be prescribed, such as orlistat, bupropion with naltrexone, phentermine with topiramate, lorcaserin, glucagon-like peptide-1 (GLP-1) receptor agonists or dual glucose insulinotropic polypeptide receptor/GLP-1 agonists (6).

For individuals struggling with severe obesity associated with comorbidities, bariatric surgery is often recommended as a solution to aid in weight loss and improve overall health. There are numerous types of bariatric surgeries, some being restrictive, others malabsorptive, and some combining both approaches. Today, the most performed type of bariatric surgery is vertical sleeve gastrectomy (VSG - restrictive) accounting for 60.4% of all the surgeries performed worldwide (7). Following VSG, where most of the stomach is surgically resected, there are deficiencies in micronutrient absorption such as iron, zinc, selenium, calcium, vitamin D, folate and B12 (8). Deficiencies in folate and vitamin B12 have previously been linked to elevated homocysteine levels (9), that can cause collagen cross-linking impairment (10), alteration of osteoclast and osteoblast activities (11, 12), leading to poor bone health and fragility fracture (13). Indeed, various studies have shown an increased risk of fractures after bariatric surgery up to a two-fold increase in fracture risk (14-17). When examining the specific sites where fracture risk is increased, it is mainly at the wrist, hip, and femoral neck (17).

As obesity is a known factor of infertility (18), there is an increasing demand of obese women of child-bearing age to undergo bariatric surgery and VSG in order to restore fertility (19). Pregnancy and lactation periods are known factors affecting skeletal health of mother with a reversible bone loss due to changes in hormonal regulation(20, 21). However little is known on the skeletal health of mother that prior to pregnancy underwent a VSG and especially whether the pregnancy/lactation periods could additively aggravate the compromised bone phenotype after VSG.

The aims of this study was therefore to develop a rat model of vertical sleeve gastrectomy followed by pregnancy and lactation periods to study their effects on skeletal health. A longitudinal follow-up by invivo microCT of the trabecular and cortical bone mass at the tibia, as well as a longitudinal evaluation of bone remodelling was performed. At sacrifice, bone pieces were further recovered and a full evaluation of bone biomechanic, mass, microarchitecture and material properties were done.

## 2. MATERIALS & METHODS

### 2.1. Animals

All procedures were carried out in accordance with the European Union Directive 2010/63/EU for animal experiments and were approved by the regional ethics committee for animal use (authorization APAFIS#24621-202003111410631). Sprague-Dawley females aged 6-week-old were fed with a high fat high sugar (HFHS) diet composed of high fat pellets (824053, Special Diet Services, Witham, UK) and sweetened condensed milk (Nestlé, Issy-les-Moulineaux, France) for 8 weeks. The supplementary figure 1 recapitulates the procedures sustained by all animal groups.

At 14 weeks of age animals were randomly allocated to one of the surgery groups represented by either sham (HFHS-Sham group, n=7), or vertical sleeve gastrectomy (HFHS-VSG group, n=10). Surgery was performed under gazous anesthesia with isoflurane supplemented with buprenorphine (0.2mg/kg bw). The VSG procedure consisted in discarding 80% of the posterior mass of the stomach as previously described (22). Three days before surgeries, animals were given vitamin supplements in drinking water (vitaRongeur, Virbac, Carros, France) for a period of 6 days. During the 72hrs post-surgery, animals received subcutaneous administrations of buprenorphine (0.2 mg/kg bw) and amoxicillin/clavulanic acid (50 mg/kg/day – 5 mg/kg/day). Eight age- and sex-matched Sprague-Dawley rats under normal rodent diet (A04, Safe, Augy, France) and sham-operated were used as controls (SD-Sham group). These animals received also the vitamin supplements given to HFHS-fed animals.

At 18 weeks of age, all animals were mated with Sprague-Dawley male rats and only operated-females that had offspring were sacrificed at 30 weeks of age. Animals were housed in social groups and maintained in a 12 hours:12 hours light:dark cycle and had free access to water and diet *ad libitum*. All animals received intraperitoneal administrations of calcein green (10 mg/kg) 10 days and 2 days before sacrifice. Bone pieces were collected at the time of sacrifice for further analyses.

### 2.2 In vivo microcomputed tomography (In vivo microCT)

A longitudinal follow-up of bone microstructure at the proximal metaphysis of the right tibia was performed by in vivo microCT under general gaseous anaesthesia before surgery, and 4,10 and 16 weeks post-surgery. Briefly, the proximal metaphysis was analysed with a Bruker 1076 microtomograph (Bruker Skyscan, Kontich) operated at 80kV and 120 μA. An isotropic voxel size of 18 μm was used with an integration time of 2000 ms and a rotation step of 0.5° using a 1-mm aluminium filter. Reconstruction of 2D projections was done using NRecon software (version 1.6.10.2) using ring artefact correction set at 10, beam hardening correction set at 25% and a median filter. For trabecular analysis, a volume of interest located 0.5 mm below the growth plate and extending 2 mm down was chosen. For cortical analysis, a volume of interest (0.5 mm) centred 3 mm below the growth plate was chosen. Global threshold set at 300 mg/cm^3^ and 700 mg/cm^3^ hydroxyapatite were used for segmenting trabecular and cortical bone tissues respectively. Between two microCT scans, regions of interest (ROI) were automatically aligned with the dataviewer software (version 1.5.6.2) using morphometrical features. Volumes of bone formation and resorption were computed with a lab-made Matlab script (Matlab R2021b, The Mathworks, Natick, MA) allowing for longitudinal evaluation of bone remodelling at the proximal metaphysis. Briefly, the two datasets were registered, and voxels present only in the older dataset were assigned to bone resorption. Voxels present only in the younger dataset were assigned to bone formation. Bone formation and bone resorption were computed as the sum of unique voxels divided by the sum of common and unique voxels. All histomorphometrical parameters were measured with the CTan software according to guidelines and nomenclature proposed by the American Society for Bone and Mineral Research.

### 2.3. Ex vivo microCT

Right tibia and femur (after 3-point bending) were analysed hydrated ex vivo by microCT with a Bruker 1272 microtomograph at 70kv, 140μA, 2000 ms integration time, an isotropic voxel size fixed at 7 μm, a rotation step set at 0.5°, and the use of a 1-mm aluminium filter. Hydroxyapatite phantoms (250 mg/cm^3^ and 750 mg/cm^3^) were used for calibration. Reconstruction of 2D projections was done using the NRecon software using ring artefact correction set at 10, beam hardening correction set at 25% and a median filter. Datasets were reaxed on the sagittal and coronal planes using the dataviewer software and transaxial sections (tibia and femur) were saved and further analysed with the CTan software. In order to analysed trabecular bone at the proximal tibia metaphysis and distal femur metaphysis, volumes of interest located 0.5 mm below/above the growth plate and extending 2 mm down/up were chosen. Trabecular bone was separated from cortical bone with an automatic contouring script in CTan software. Cortical analysis was restricted to tibia and femur, and the analyzed ROI (0.5 mm in length) was centered at 3mm under/above the growth plate. Bone was segmented from soft tissue using global thresholding set at 300 mg/cm^3^ for trabecular bone and 700 mg/cm^3^ for cortical bone. All histomorphometrical parameters were measured with the CTan software according to guidelines and nomenclature proposed by the American Society for Bone and Mineral Research.

### 2.4 Bone biomechanical test

Bending strength was measured by 3-point bending as described previously (23) and in accordance with published guidelines (24) using a constant span length of 15 mm. Right femurs were tested in the anteroposterior axis with the posterior surface facing upward, centred on the support, and the pressing force was applied vertically to the midshaft of the bone. Each bone was tested fully hydrated and at room temperature with a loading speed of 0.5 mm.min^−1^ until failure with a 500-N load cell on an Instron 5942 device (Instron, Elancourt, France), and the load–displacement curve was recorded at a rate of 100 Hz using Bluehill 3 software (Instron). Ultimate load, ultimate displacement, stiffness, and work to fracture were calculated as indicated in Turner and Burr (25). The yield load was determined as the point at which a regression line that represents a 10% loss in stiffness crossed the load–displacement curve. Postyield displacement was computed as the displacement between yielding and fracture. Additionally, tissue strength was calculated as the peak moment bending divided by the section modulus as reported in (26).

### 2.5. Raman spectroscopy

After 3-point bending, the proximal part of the right femur was embedded undecalcified in polymethymethacrylate (pMMA) at 4°C as previously reported (27). Thick bone sections (1 mm-thickness) were cut at the diaphyseal side, and the section surface was then grinded with sandpaper and polished with diamond paste (Struers, Champ sur Marne, France). In order to account for tissue age by using double calcein labellings, sections were imaged with a laser scanning confocal microscope (Leica DM8, Leica microsystems, Nanterre) set with an excitation light at 488 nm and an emission range 515-550 nm. Spectra were acquired using a Renishaw inVia Qontor confocal Raman microscope equipped with a 785 nm line laser, and a charged-coupled device (CCD) array detector. The laser power was set at 30 mW. The laser beam was focused onto the sample surface through an objective of 20 X and 0.4 numerical aperture, providing a spatial resolution of 2.4 μm and limiting polarization artefacts. Prior to each scan, spectral calibration was done using an internal silicon standard. The spectral range was set between 800 to 1800 cm^−1^, the acquisition time at 20s and 3 accumulations. All data acquisition was performed using Renishaw WiRE v.5.5 software. Spectral artefacts including cosmic rays and background were removed using the nearest neighbour method and polynomial fit, respectively. Contribution of the embedding resin was subtracted using the major peak located at ∼812 cm^-1^. Bone material parameters were: mineral-to-matrix ratio (intensity ratio between v1PO_4_ at ∼960 cm^-1^ and Amide I at ∼1668 cm^-1^), carbonate/phosphate ratio (intensity v3CO_3_ at ∼1070 cm^-1^ and v1PO_4_), nanoporosity, computed before pMMA subtraction (intensity ratio between pMMA at ∼812 cm^-1^ and v1PO_4_), mineral crystallinity (as the inverse of full-width at half maximum of the v1PO_4_), proline hydroxylation content (intensity ratio between hydroxyproline at ∼872 cm^-1^ and proline backbone at ∼854 cm^-1^), glycosaminoglycan content (intensity ratio between GAG at ∼1375 cm^-1^ and Amide III at ∼1245 cm^-1^), collagen maturity (intensity ratio between ∼1670 cm^-1^ and ∼1690 cm^-1^).

Spectra were acquired in two different locations : (I) to investigate bone material properties at site of bone formation, spectra were randomly positioned between double calcein labellings and acquired as indicated above, and (II) to investigate bone material properties on the full cortical width, a line of points was drawn on the anterior quadrant extending from the periosteal to endosteal surfaces with a spatial resolution of 6 μm and acquired as above. For the later, histogram distribution for each compositional parameters were fitted with a Gaussian model and considered normally distributed if the R^2^ coefficient was >0.95. In the present study, no histogram deviated from normal distribution. For each of the compositional parameters, the mean of the pixel distribution (excluding non-bone pixels) was computed as:

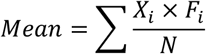

With Xi, representing the value of the compositional parameter for the i^th^ bin, Fi, representing the percentage of bone area with the Xi value and N, representing the total number of bins in the histogram distribution. The full width of the half maximum of each distribution was also computed to determine heterogeneity in the compositional parameter.

### 2.6 Statistical analysis

Statistical analyses were performed with GraphPad Prism 8.0 (GraphPad Software, La Jolla, CA, USA). The Brown-Forsythe test of equal variance was applied, and when normality was respected, ordinary one-way ANOVA with multiple post hoc Dunnett comparisons were conducted. When normality was not respected, Kruskal-Wallis tests with multiple post hoc Dunn’s comparisons were carried out. A linear regression was calculated to establish a correlation between the results of the in vivo microCT study and those of the ex vivo microCT study. Differences at *p* < 0.05 were considered significant.

## 3. RESULTS

### 3.1. Longitudinal evaluation of bone mass and remodelling at the proximal tibia metaphysis

The right tibia of all animals was scanned just before surgery, 4 weeks after surgery, 10 weeks post-surgery, corresponding to weaning, and 16 weeks post-surgery at sacrifice. Interestingly, in the SD+Sham group, animals sustained a trabecular bone loss by ∼-54% (p<0.001) at weaning but quickly gained trabecular bone mass post-weaning to reach a ∼66% (p<0.001) augmentation in this parameter at sacrifice (Figure 1). These fluctuations of bone mass were accompanied by modifications of the pattern of bone formation with a significant decrease in trabecular bone formation by 9% (p<0.001) at weaning and a significant increase by 10% at sacrifice. No significant modification of the pattern of bone resorption was observed in this group. In the HFHS-Sham group, a significant 64% decrease (p<0.001) in BV/TV was observed at weaning and accompanied by significant reduction in bone formation (-6%, p=0.017) and augmentation in bone resorption (+33%, p<0.001). At sacrifice, all HFHS-Sham animals gained BV/TV by ∼83% (p<0.001) due to increase in bone formation (+26%, p=0.023) and decrease in bone resorption (-21%, p=0.024). In the HFHS-VSG group, in opposition to HFHS-Sham and SD-Sham, a significant loss of BV/TV by approximately 55% (p<0.001) was evidenced 4 weeks post-surgery. Interestingly, these animals did not lose further bone at weaning (p=0.50). However, during the weaning-sacrifice period, these animals presented with the largest gain in BV/TV (+75%, p<0.001) associated with a 19% (p=0.003) augmentation in bone formation and a 22% (p=0.047) reduction in bone resorption.

**Figure 1:**
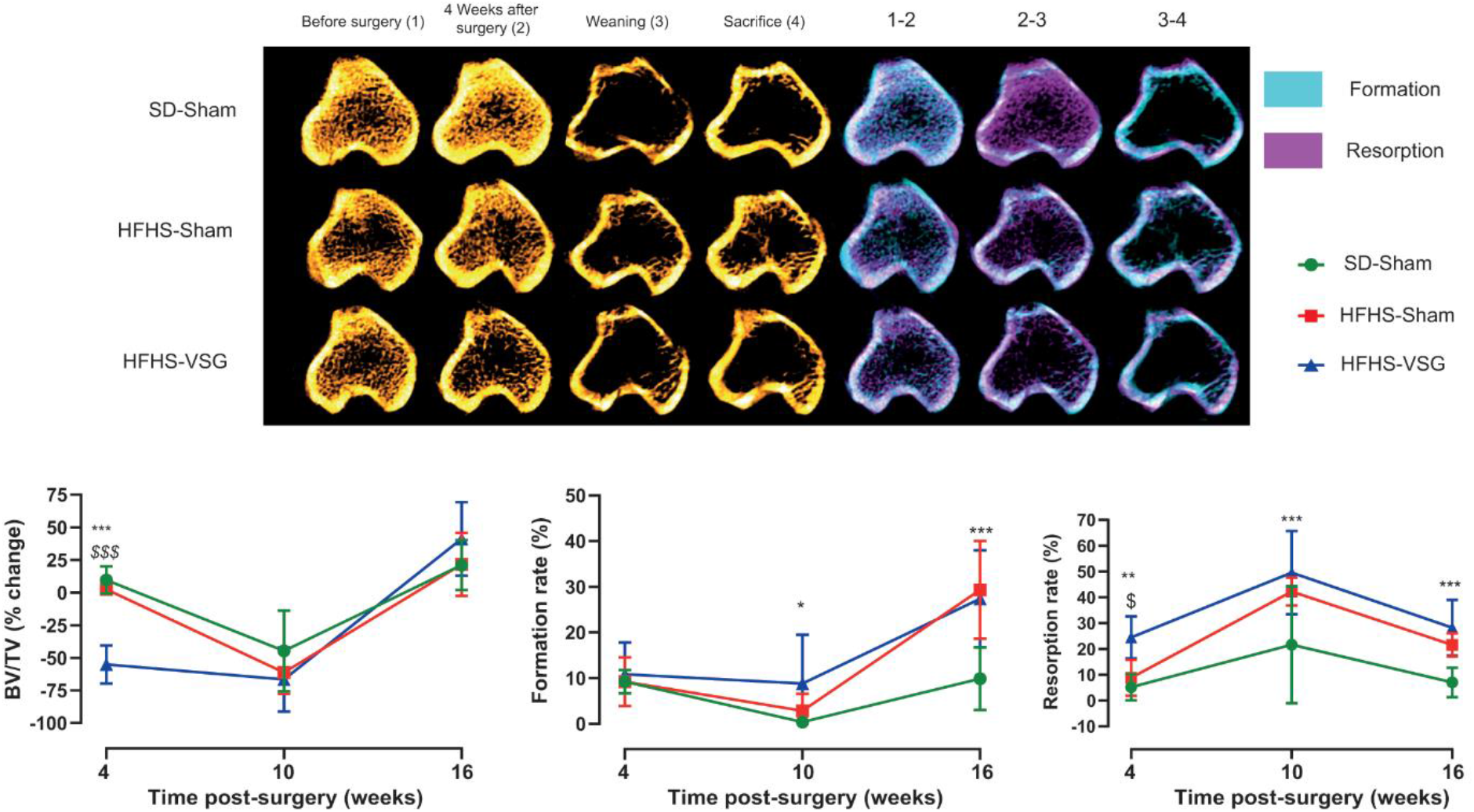
In vivo follow-up investigation of trabecular bone mass and remodelling at the proximal tibia metaphysis. MicroCT images taken before surgery (1), 4 weeks after surgery (2), at weaning (3) and at sacrifice (4). Bone formation and resorption rates were computed before and 4 weeks after surgery (period 1-2), between 4 weeks post-surgery and weaning (period 2-3), and between weaning and sacrifice (period 3-4). Quantitative analysis of BV/TV is reported as % change from previous time point and not from baseline. Statistical differences between SD-Sham (n=8), HFHS-Sham (n=7) and HFHS-VSG (n=10) were computed using a one-way ANOVA with Tukey multiple comparisons. *: p<0.05; **:p<0.001; ***:p<0.0001 vs. SD-Sham and^$^: p<0.05; ^$$$^:p<0.0001 vs. HFHS-Sham at each time point.

### 3.2. Longitudinal evaluation of bone mass and remodelling at the tibia midshaft

At the tibia midshaft, in the SD-Sham group, a significant augmentation (+18%, p=0.037) of cortical bone mass, represented by Ct.Ar/Tt.Ar, was evidenced between pre- and post-surgery evaluations (Figure 2). Ct.Ar/Tt.Ar was not further modified significantly at weaning but was augmented by 38% (p<0.001) at sacrifice and associated with significant cortical resorption (-9%, p=0.029). In the HFHS-Sham group, no significant alteration of Ct.Ar/Tt.Ar was evidenced between pre- and post-surgery period. However, at weaning these animals presented with a significant reduction in Ct.Ar/Tt.Ar (-14%, p=0.016). At sacrifice, Ct.Ar/Tt.Ar was augmented by 47% (p=0.005), as compared with weaning, and associated with significant augmentation of cortical formation (17%, p=0.025), and reduction in cortical resorption (12%, p=0.006). In the HFHS-VSG group, animals reduced significantly Ct.Ar/Tt.Ar by 13% (p=0.002). At weaning, these animals did not present with a significant alteration of Ct.Ar/Tt.Ar (p=0.37). However, at sacrifice, Ct.Ar/Tt.Ar was significantly augmented by 35% (p=0.033) and associated with a significant reduction in cortical resorption (-22%, p=0.027).

**Figure 2:**
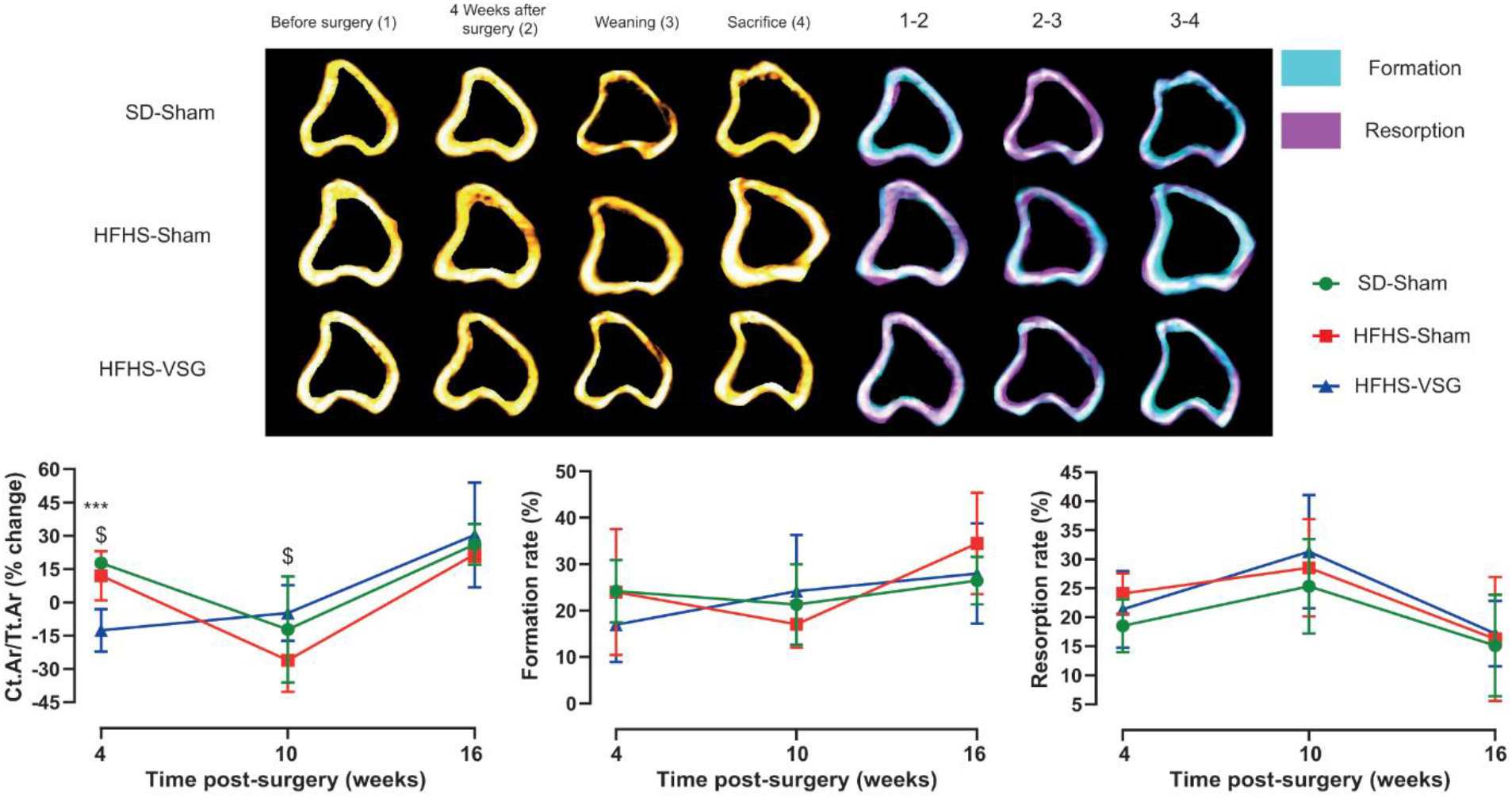
In vivo follow-up investigation of cortical bone mass and remodelling at the tibia midshaft. MicroCT images taken before surgery (1), 4 weeks after surgery (2), at weaning (3) and at sacrifice (4). Bone formation and resorption rates were computed before and 4 weeks after surgery (period 1-2), between 4 weeks post-surgery and weaning (period 2-3), and between weaning and sacrifice (period 3-4). Quantitative analysis of Ct.Ar/Tt.Ar is reported as % change from previous time point and not from baseline. Statistical differences between SD-Sham (n=8), HFHS-Sham (n=7) and HFHS-VSG (n=10) were computed using a one-way ANOVA with Tukey multiple comparisons. ***:p<0.0001 vs. SD-Sham and ^$^: p<0.05 vs. HFHS-Sham at each time point.

### 3.3. Ex-vivo evaluation of trabecular microarchitecture at the proximal tibia metaphysis and cortical microarchitecture at the tibia midshaft

After sacrifice, the tibia microarchitecture was analysed with a higher resolution as used for longitudinal evaluation of bone mass and remodelling. Figure 3A represents 3D-models of each animal group. As evidenced in Figure 3B, trabecular bone mineral densities (BMD) were significantly reduced in HFHS-Sham (-35%) and HFHS-VSG (-39%) as compared with SD-Sham, respectively. However, no significant differences were noted between the two HFHS groups. These reductions of BMD were mirrored by reductions in BV/TV in HFHS-Sham (-36%) and HFHS-VSG (-42%) as compared with SD-Sham controls. Decreases in Tb.N, but not Tb.Th, were evidenced in HFHS-Sham (-34%) and HFHS-VSG (-41%) as compared with SD-Sham. Interestingly, the ex vivo BV/TV was significantly correlated with the BV/TV measured from the longitudinal assessment at sacrifice (R^2^ = 0.42, p<0.001), validating the longitudinal follow-up.

**Figure 3:**
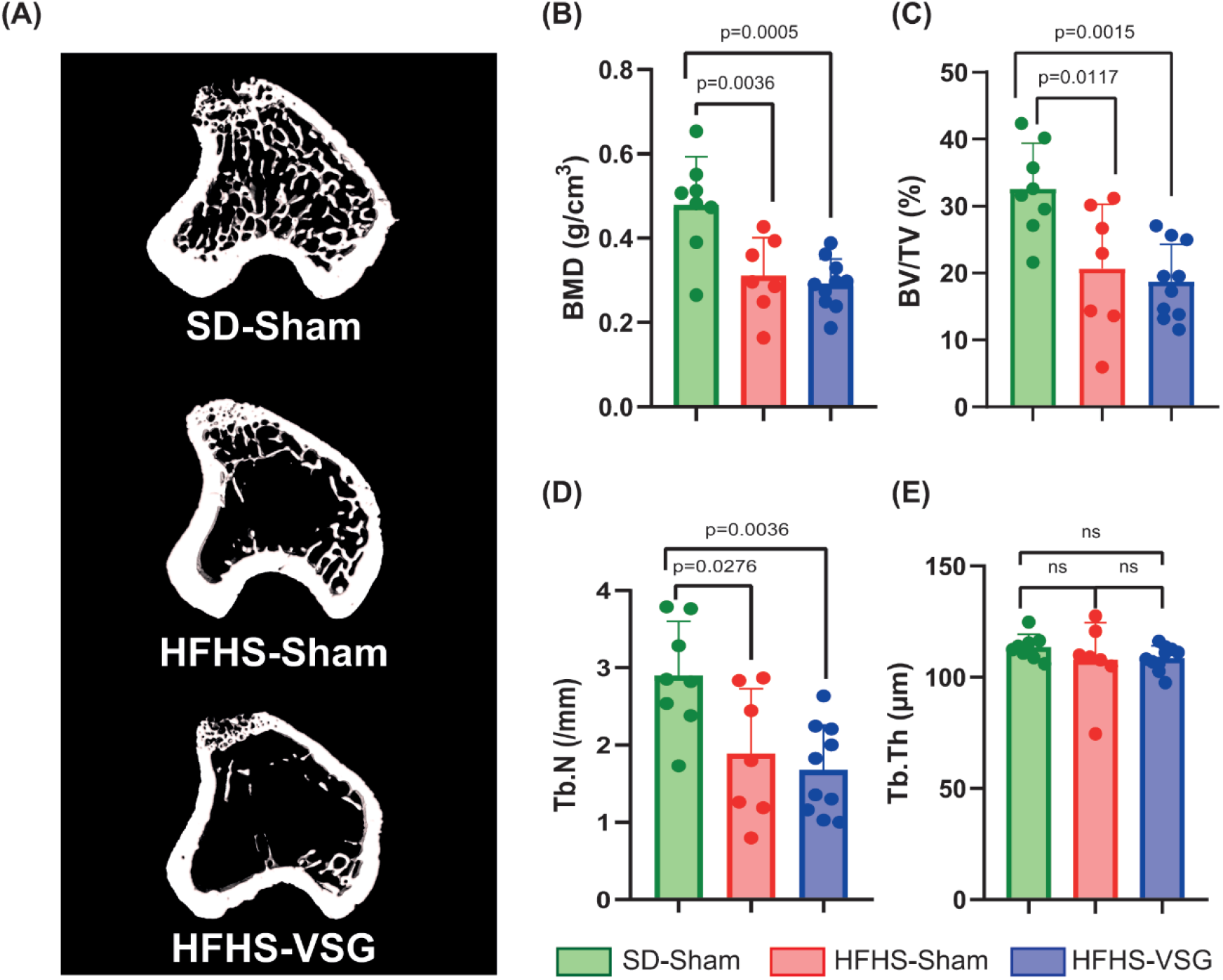
Ex vivo assessment of trabecular bone mass and microarchitecture at the proximal tibia metaphysis. (A) 3D model of each animal group. (B-E) BMD, BV/TV, Tb.N, Tb.Th were measured by microCT. Statistical differences between SD-Sham (n=8), HFHS-Sham (n=7) and HFHS-VSG (n=10) were computed using a one-way ANOVA with Tukey multiple comparisons.

At the tibia midshaft, 3D models did not suggest a specific alteration of cortical microarchitecture (Figure 4) and indeed, none of the conventional cortical microarchitecture parameters was significantly altered by HFHS or Sham/VSG surgeries. However, and in parallel to what was seen in the proximal metaphysis, Ct.Ar/Tt.Ar measured ex vivo was significantly correlated to the measure at sacrifice on the longitudinal evaluation (R^2^ = 0.42, p=0.001).

**Figure 4:**
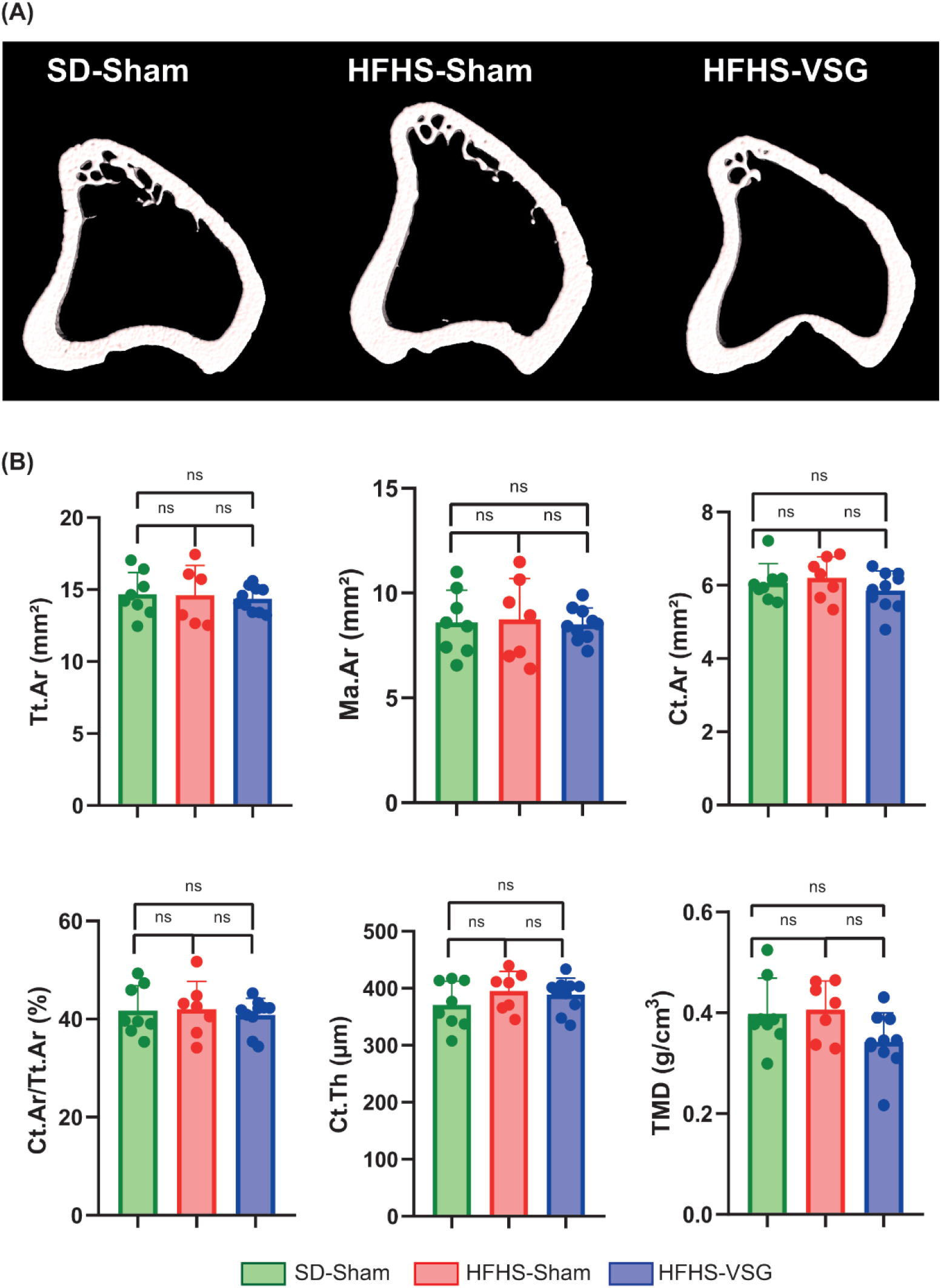
Ex vivo assessment of cortical microarchitecture at the tibia midshaft. (A) 3D model of each animal group. (B) Tt.Ar, Ma.Ar, Ct.Ar, Ct.Ar/Tt.Ar, Ct.th and TMD were measured by microCT. Statistical differences between SD-Sham (n=8), HFHS-Sham (n=7) and HFHS+VSG (n=10) were computed using a one-way ANOVA with Tukey multiple comparisons.

### 3.4 Evaluation of biomechanical response

As bariatric surgery is linked to excessive bone fragility, we performed 3-point bending measurement at the femur mid-diaphysis in all three groups. Animals under HFHS diet, albeit of the type of surgery, exhibited significant lower ultimate load (20% - 26%), yield load (30 - 32%), stiffness (19% - 23%) and work-to-fracture (18% - 20%) as compared with SD-Sham animals (Table 1). In order to ascertain whether changes in bone biomechanics were due to bone microarchitecture or changes in bone matrix biomechanics, we also computed tissue strength. Interestingly, tissue strength was significantly lower in HFHS-Sham (-21%, p=0.008) and HFHS-VSG (-21%, p=0.005) animals as compared with SD-Sham suggesting a possible alteration of bone material properties.

**Table 1:**
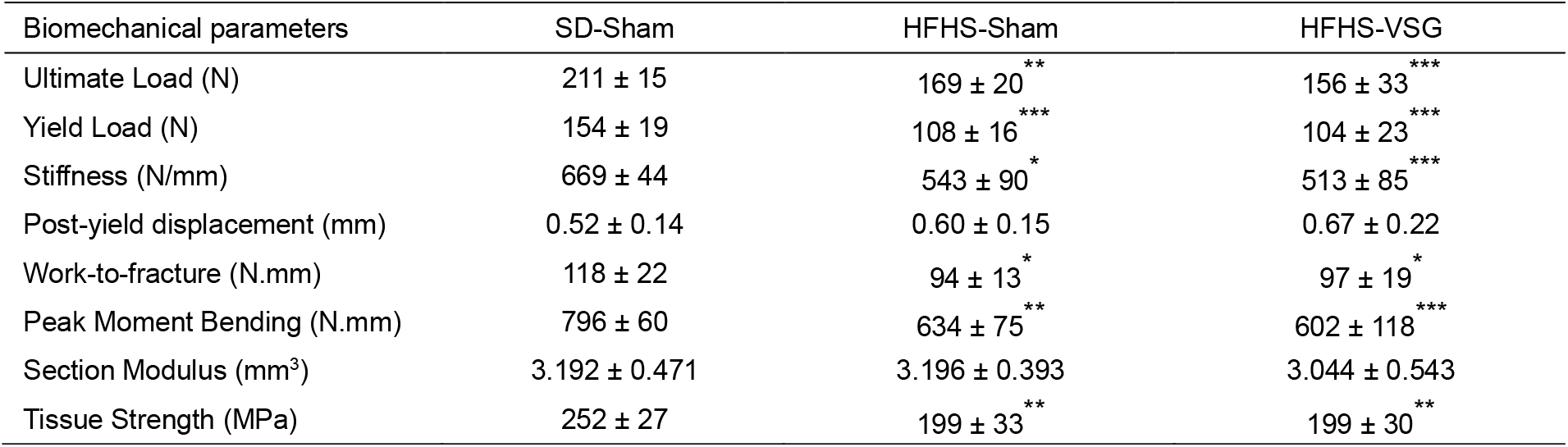
Biomechanical parameters measured by 3-point bending at the midshaft femur. Statistical differences between SD-Sham (n=8), HFHS-Sham (n=7) and HFHS-VSG (n=10) were investigated by one-way ANOVA with Tukey’s multiple comparison tests. ^*^: p<0.05, ^**^: p<0.001, ^***^: p<0.0001 vs. SD-Sham.

### 3.5. Evaluation of femur microarchitecture and geometry

In order to ascertain whether alterations of bone biomechanics was due to specific alterations of bone microstructure at the femur that could not be observed at the tibia, we evaluated trabecular BMD and BV/TV at the distal femoral metaphysis (Figure 5). Similar alterations as observed at the proximal tibia metaphysis were encountered and represented by lower BMD (-42% and -55%, p=0.0068 and p=0.0003) and BV/TV (-42% and -52%, p=0.0016 and p<0.0001) in HFHS-Sham and VSG groups, respectively. At the femoral diaphysis, as evidenced already at the tibia, no significant alterations of cortical microarchitecture could be evidenced between groups. However, tissue mineral density (TMD) was significantly lower in HFHS groups (-25% and -25%, p=0.045 and p=0.041) as compared with SD-Sham. No alterations of femur geometry were observed between groups. On the other hand, TMD was positively correlated with yield load (R^2^=0.20, p=0.028).

**Figure 5:**
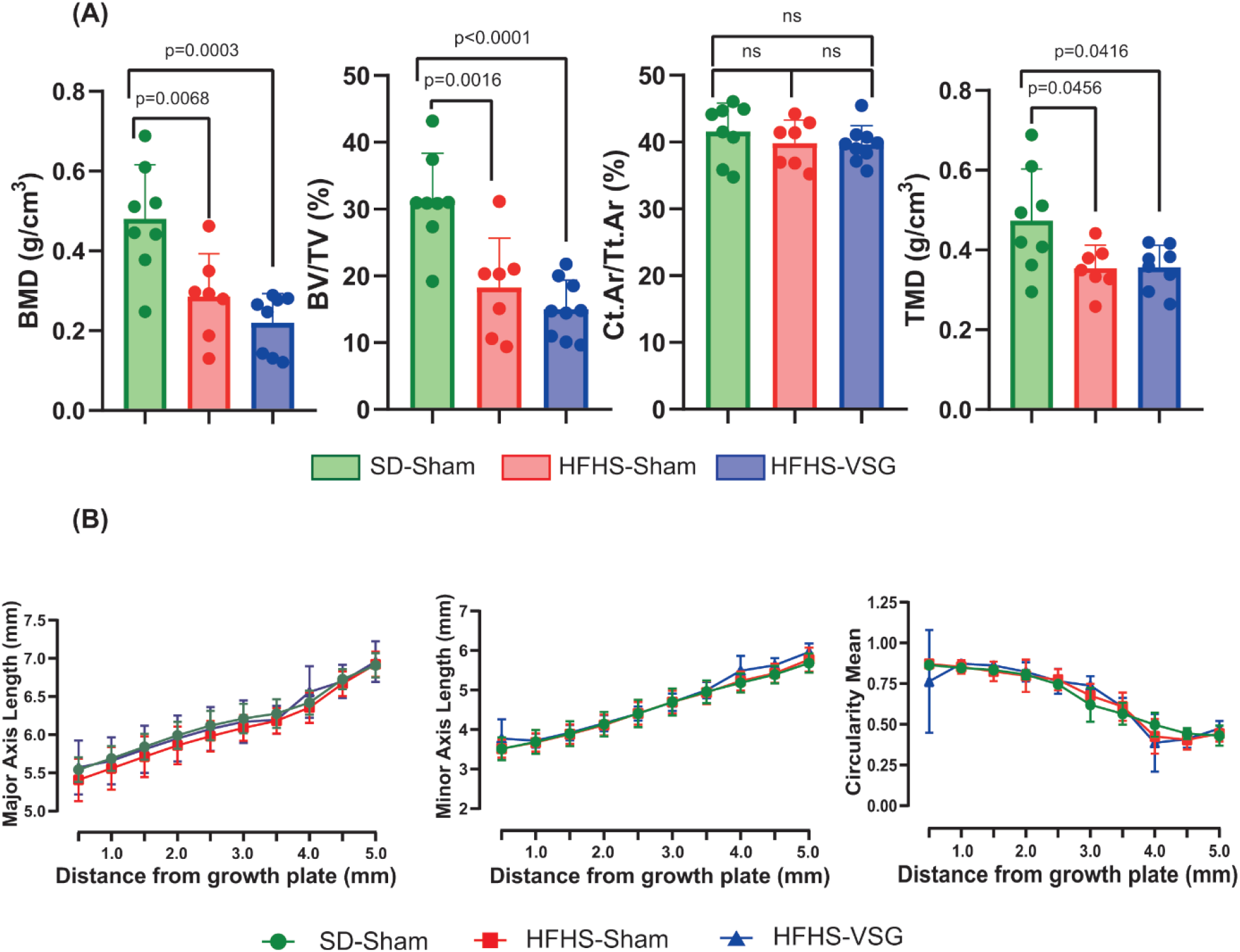
Ex vivo assessment of trabecular and cortical bone microarchitecture at the distal femur metaphysis and femur midshaft. (A) BMD, BV/TV, Ct.Ar/Tt.Ar and TMD were obtained by microCT. (B) Major axis length, minor axis length and circularity mean were measured from microCT scan. Statistical differences between SD-Sham (n=8), HFHS-Sham (n=7) and HFHS-VSG (n=10) were computed using a one-way ANOVA with Tukey multiple comparisons

### 3.6. Evaluation of bone material properties at the femur mid-diaphysis

We then investigated the extracellular matrix properties by Raman spectroscopy on transaxial section of the mid-femur (Table 2). At the bone formation site, no significant differences between all three groups of animals were encountered. Interestingly, no alterations of the v1PO_4_/Amide I ratio, indicative of the degree of mineralization, was evidenced. We also examined material properties of the bone matrix by performing Raman spectroscopy over the full cortical width. In all animals, each parameter followed a gaussian distribution. As such the mean and full width at half maximum (FWHM) were computed. Matrix nanoporosity was significantly lower in HFHS-VSG animals (-50%, p=0.040) as compared to HFHS-Sham Interestingly, the FWHM of v1PO_4_/Amide I was significantly lowered in HFHS-Sham (-53%, p=0.035) and HFHS-VSG (-54%, p=0.012) as compared with SD-Sham.

**Table 2:** Bone material properties of the femur midshaft measured by Raman microspectroscopy. Bone material properties were measured either at bone formation site by placing 3 points between double calcein labels or by drawing a line over the full cortical width. The latter allows for computation of parameter distribution, represented by a gaussian shape, and reported as mean and full width at half maximum (FWHM). Statistical differences between SD-Sham (n=8), HFHS-Sham (n=7) and HFHS-VSG (n=10) were computed using a one-way ANOVA with Tukey multiple comparisons. *: p<0.05 vs. SD-Sham and ^#^: p<0.05 vs. HFHS-Sham.

| Bone material parameters | SD-Sham | HFHS-Sham | HFHS-VSG |
| --- | --- | --- | --- |
| **At bone formation site** |  |  |  |
| $\nu_1\text{PO}_4/\text{Amide I}$ | $10.83 \pm 2.9$ | $11.83 \pm 2.7$ | $10.02 \pm 3.1$ |
| $\nu_3\text{CO}_3/\nu_1\text{PO}_4$ | $0.164 \pm 0.006$ | $0.165 \pm 0.009$ | $0.162 \pm 0.010$ |
| Nanoporosity | $73 \pm 30.3$ | $61.6 \pm 44.4$ | $121.1 \pm 85.4$ |
| Mineral crystallinity | $0.106 \pm 0.01$ | $0.110 \pm 0.008$ | $0.108 \pm 0.009$ |
| Proline hydroxylation content | $0.62 \pm 0.047$ | $0.64 \pm 0.05$ | $0.69 \pm 0.08$ |
| GAG/Amide III | $0.0863 \pm 0.006$ | $0.0936 \pm 0.016$ | $0.0053 \pm 0.015$ |
| $1670 \text{ cm}^{-1}/1690 \text{ cm}^{-1}$ | $2.739 \pm 0.59$ | $2.996 \pm 0.76$ | $2.233 \pm 0.37$ |
| **At full cortical width** |  |  |  |
| Mean $\nu_1\text{PO}_4/\text{Amide I}$ | $11.18 \pm 1.0$ | $11.93 \pm 0.75$ | $11.88 \pm 1.11$ |
| Mean $\nu_3\text{CO}_3/\nu_1\text{PO}_4$ | $0.183 \pm 0.004$ | $0.178 \pm 0.003$ | $0.180 \pm 0.005$ |
| Mean Nanoporosity | $0.0037 \pm 0.0015$ | $0.0042 \pm 0.0027$ | $0.0020 \pm 0.0007^\#$ |
| Mean Mineral crystallinity | $0.055 \pm 0.0002$ | $0.055 \pm 0.0002$ | $0.0550 \pm 0.0003$ |
| Mean Proline hydroxylation content | $0.76 \pm 0.02$ | $0.74 \pm 0.01$ | $0.75 \pm 0.02$ |
| Mean GAG/Amide III | $0.475 \pm 0.015$ | $0.443 \pm 0.014$ | $0.454 \pm 0.014$ |
| Mean $1670 \text{ cm}^{-1}/1690 \text{ cm}^{-1}$ | $2.773 \pm 0.079$ | $2.802 \pm 0.25$ | $2.732 \pm 0.093$ |
| FWHM $\nu_1\text{PO}_4/\text{Amide I}$ | $4.1 \pm 2.1$ | $2.0 \pm 0.7^*$ | $1.9 \pm 0.6^*$ |
| FWHM $\nu_3\text{CO}_3/\nu_1\text{PO}_4$ | $0.014 \pm 0.002$ | $0.012 \pm 0.006$ | $0.012 \pm 0.006$ |
| FWHM Nanoporosity | $0.003 \pm 0.001$ | $0.003 \pm 0.001$ | $0.003 \pm 0.001$ |
| FWHM Mineral crystallinity | $0.0014 \pm 0.0005$ | $0.0013 \pm 0.0003$ | $0.0013 \pm 0.0002$ |
| FWHM Proline hydroxylation content | $0.15 \pm 0.04$ | $0.17 \pm 0.01$ | $0.13 \pm 0.05$ |
| FWHM GAG/Amide III | $0.123 \pm 0.017$ | $0.137 \pm 0.008$ | $0.125 \pm 0.021$ |
| FWHM $1670 \text{ cm}^{-1}/1690 \text{ cm}^{-1}$ | $0.2290 \pm 0.017$ | $0.2148 \pm 0.006$ | $0.2177 \pm 0.006$ |

The FWHM of v1PO_4_/Amide I was positively correlated with the peak moment bending (R^2^=0.22, p=0.022). On the other hand, nanoporosity, that was significantly altered in HFHS-VSG as compared with HFHS-Sham, was not correlated with peak moment bending (R^2^=0.03, p=0.48).

## 4. DISCUSSION

Bariatric surgery has been associated with an increased bone fragility with a mechanism that remains to be fully elucidated. As bariatric surgery and vertical sleeve gastrectomy is increasingly performed in young obese female in order to restore fertility, we thought to ascertain whether a gestation followed by a lactation period could affect the bone fragility observed after VSG. In this study, we evidenced, as others (28-31), that vertical sleeve gastrectomy resulted in rapid trabecular and cortical bone loss at the appendicular skeleton due to a significant increase in bone resorption. However, the gestation/lactation period seems to revert the bone phenotype with an increase in trabecular and cortical bone mass from end of lactation period until sacrifice. Interestingly, we also evidenced that bone fragility was present in HFHS animals independently of the type of surgery (Sham or VSG) and associated to lower tissue strength and alterations of mineralization heterogeneity. However, matrix nanoporosity was only affected in HFHS-VSG. This suggested that the diet and perturbations of lipid and carbohydrate metabolisms might be responsible of the bone fragility and alterations of material properties rather than the VSG surgery per se.

The bone loss due to calcium mobilization during gestation and lactation is more significant in a rodent model because litter size is larger. In rats, there is typically a reduction in bone mass of 15 to 35% at the spine and hip respectively, accompanied by a decrease in trabecular and cortical thickness and a reduction in the number of bone trabeculae (32). Nevertheless, whether in female individuals or rodents, and even though the exact mechanisms are not fully understood, an increase in osteoclast resorption during gestation/lactation period followed by an increase in the population of active osteoblasts at the end of lactation has been highlighted (33). This could explain the recovery of trabecular bone mobilized during lactation and observed in our study, especially with respect to the higher bone formation observed in the longitudinal follow-up. Although not investigated in the present study, the augmentation in active osteoblasts following the lactation period may involves several endocrine mediators such as estradiol, prolactin, vitamin D, parathyroid hormone, parathyroid hormone-related peptide, calcitonin, and gonadotrophins (32-35).

Obesity is associated with increases in BMI and BMD (36) despite a higher risk of fractures in obese children, adolescents, and adults (3, 37). It is important to ascertain whether rodent models of obesity could recapitulate the findings observed in humans. Several animal studies have showed that a high-fat diet leads to bone repercussions, including a decrease in BV/TV and Tb.N in long bones (38, 39) despite higher body weight. As such the bone phenotype in rodent is not identical as the one seen in humans. Nevertheless, in the present study, we reported alterations of trabecular bone mass in animals under a HFHS diet and after pregnancy/lactation periods. Interestingly, although fertility and energy metabolism have been ameliorated following bariatric surgery (40-45),it seems that bone alterations due to HFHS diet are unaffected by vertical sleeve gastrectomy.

On the other hand, and as hypothesized as the beginning of this study, it seems that pregnancy and lactation did not aggravate the bone phenotype due to VSG. More importantly, changes in patterns of bone resorption and bone formation following these periods appeared similar between HFHS-Sham and HFHS-VSG suggesting that bariatric surgery is not detrimental to bone remodeling programming. However, despite similar cortical geometry and microarchitecture at the femur, HFHS animals exhibited a bone fragility that seems to be associated with the quality of the extracellular matrix, especially with the lack of heterogeneity in the degree of mineralization. This alterations has previously been observed in atypical fracture in women with bisphosphonate treatment (46, 47). A plausible scenario is represented by adipokine contribution. Indeed, adipose tissue is known to release pro-inflammatory cytokines (IL-6, TNF-α, IL-1β) that reduce osteoblastic differentiation of mesenchymal stem cells, decrease bone formation (48) allowing more time for the matrix to mineralised and reducing heterogeneity.

A strength of our study is represented by the longitudinal follow-up of bone mass and bone remodeling that allowed us to precisely quantify 3D bone remodeling and trabecular and cortical microarchitecture at different time points, which, to our knowledge, has never been done in a rat model before. Without this follow-up, we would have overlooked the bone loss induced by VSG, as animals on the HFHS diet appear identical at terminal endpoints in terms of trabecular and cortical bone. Another strength relies in the full assessment of trabecular and cortical bone mass, microarchitecture, biomechanics and material properties that allow to evidenced that alteration of bone biomechanics is due to changes in matrix properties rather than changes in bone microarchitecture. A limitation of this study is represented by the model organism that does not fully reproduced the bone phenotype observed in human individuals, especially in term of higher BMD due to higher body weight and the structure of the cortical bone that is not haversian in rodents. Nevertheless, similar trabecular bone loss at the tibia proximal metaphysis were observed rapidly after VSG, similar to what is observed after bariatric surgery in human individuals (49).

## 5. CONCLUSIONS

In our study, we demonstrated that pregnancy and lactation periods after vertical sleeve gastrectomy did not aggravate the bone fragility in a rodent model. However, this study suggests that bone fragility due to high fat high sugar diet, represented by modifications of material properties, is not reversed by vertical sleeve gastrectomy, at least at cortical bone site. This requires further investigation in humans to ascertain whether bariatric surgery can revert obesity-induced bone fragility.

## 6. AUTHOR CONTRIBUTION

**Malory Couchot:** Investigation, Formal analysis, Writing - Original Draft; **Francoise Schmitt:** Conceptualization, Investigation, Writing – Review & Editing; **Morgane Mermet:** Investigation, Writing - Review & Editing; **Celine Fassot:** Conceptualization, Resources, Writing - Review & Editing; **Guillaume Mabilleau:** Conceptualization, Formal analysis, Writing - Review & Editing, Supervision, Funding acquisition, Data curation

## 7. ACKNOWLEDGEMENTS

The authors thank the HiMolA Platform, University of Angers for assistance with bone evaluation. The authors are grateful to the SCAHU platform, SFR ICAT, University of Angers for animal housing and care. Part of this work was supported by a grant from Fondation de l’Avenir (grant number AP-RM-20-026).

## 9. DATA AVAILABILITY STATEMENT

The data that support the findings of this study are available from the corresponding author upon reasonable request.

## REFERENCES

1. Chen R, Armamento-Villareal R. Obesity and Skeletal Fragility. J Clin Endocrinol Metab. 2024;109(2):e466–e77.

2. WHO. Obesity and overweight 2024 [Available from: https://www.who.int/news-room/fact-sheets/detail/obesity-and-overweight.

3. Mosca LN, Goldberg TB, da Silva VN, da Silva CC, Kurokawa CS, Bisi Rizzo AC, et al. Excess body fat negatively affects bone mass in adolescents. Nutrition. 2014;30(7-8):847–52.

4. Rinonapoli G, Pace V, Ruggiero C, Ceccarini P, Bisaccia M, Meccariello L, et al. Obesity and Bone: A Complex Relationship. Int J Mol Sci. 2021;22(24).

5. Elmaleh-Sachs A, Schwartz JL, Bramante CT, Nicklas JM, Gudzune KA, Jay M. Obesity Management in Adults: A Review. JAMA. 2023;330(20):2000–15.

6. Kosmalski M, Deska K, Bak B, Rozycka-Kosmalska M, Pietras T. Pharmacological Support for the Treatment of Obesity-Present and Future. Healthcare (Basel). 2023;11(3).

7. Kamal FA, Fernet LY, Rodriguez M, Kamal F, Da Silva NK, Kamal OA, et al. Nutritional Deficiencies Before and After Bariatric Surgery in Low- and High-Income Countries: Prevention and Treatment. Cureus. 2024;16(2):e55062.

8. Mechanick JI, Apovian C, Brethauer S, Garvey WT, Joffe AM, Kim J, et al. Clinical Practice Guidelines for the Perioperative Nutrition, Metabolic, and Nonsurgical Support of Patients Undergoing Bariatric Procedures - 2019 Update: Cosponsored by American Association of Clinical Endocrinologists/American College of sEndocrinology, the Obesity Society, American Society for Metabolic & Bariatric Surgery, Obesity Medicine Association, and American Society of Anesthesiologists - Executive Summary. Endocr Pract. 2019;25(12):1346–59.

9. Selhub J, Jacques PF, Wilson PW, Rush D, Rosenberg IH. Vitamin status and intake as primary determinants of homocysteinemia in an elderly population. JAMA. 1993;270(22):2693–8.

10. Liu G, Nellaiappan K, Kagan HM. Irreversible inhibition of lysyl oxidase by homocysteine thiolactone and its selenium and oxygen analogues. Implications for homocystinuria. J Biol Chem. 1997;272(51):32370–7.

11. Herrmann M, Widmann T, Colaianni G, Colucci S, Zallone A, Herrmann W. Increased osteoclast activity in the presence of increased homocysteine concentrations. Clin Chem. 2005;51(12):2348–53.

12. Herrmann M, Umanskaya N, Wildemann B, Colaianni G, Widmann T, Zallone A, et al. Stimulation of osteoblast activity by homocysteine. J Cell Mol Med. 2008;12(4):1205–10.

13. van Meurs JB, Dhonukshe-Rutten RA, Pluijm SM, van der Klift M, de Jonge R, Lindemans J, et al. Homocysteine levels and the risk of osteoporotic fracture. N Engl J Med. 2004;350(20):2033–41.

14. Douglas IJ, Bhaskaran K, Batterham RL, Smeeth L. Bariatric Surgery in the United Kingdom: A Cohort Study of Weight Loss and Clinical Outcomes in Routine Clinical Care. PLoS Med. 2015;12(12):e1001925.

15. Fashandi AZ, Mehaffey JH, Hawkins RB, Schirmer B, Hallowell PT. Bariatric surgery increases risk of bone fracture. Surg Endosc. 2018;32(6):2650–5.

16. Lu CW, Chang YK, Chang HH, Kuo CS, Huang CT, Hsu CC, et al. Fracture Risk After Bariatric Surgery: A 12-Year Nationwide Cohort Study. Medicine (Baltimore). 2015;94(48):e2087.

17. Rousseau C, Jean S, Gamache P, Lebel S, Mac-Way F, Biertho L, et al. Change in fracture risk and fracture pattern after bariatric surgery: nested case-control study. BMJ. 2016;354:i3794.

18. Marinelli S, Napoletano G, Straccamore M, Basile G. Female obesity and infertility: outcomes and regulatory guidance. Acta Biomed. 2022;93(4):e2022278.

19. Young MT, Phelan MJ, Nguyen NT. A Decade Analysis of Trends and Outcomes of Male vs Female Patients Who Underwent Bariatric Surgery. J Am Coll Surg. 2016;222(3):226–31.

20. Ardeshirpour L, Brian S, Dann P, VanHouten J, Wysolmerski J. Increased PTHrP and decreased estrogens alter bone turnover but do not reproduce the full effects of lactation on the skeleton. Endocrinology. 2010;151(12):5591–601.

21. Wysolmerski JJ. Interactions between breast, bone, and brain regulate mineral and skeletal metabolism during lactation. Ann N Y Acad Sci. 2010;1192:161–9.

22. Ayer A, Borel F, Moreau F, Prieur X, Neunlist M, Cariou B, et al. Techniques of Sleeve Gastrectomy and Modified Roux-en-Y Gastric Bypass in Mice. J Vis Exp. 2017(121).

23. Chretien A, Couchot M, Mabilleau G, Behets C. Biomechanical, Microstructural and Material Properties of Tendon and Bone in the Young Oim Mice Model of Osteogenesis Imperfecta. Int J Mol Sci. 2022;23(17).

24. Jepsen KJ, Silva MJ, Vashishth D, Guo XE, van der Meulen MC. Establishing biomechanical mechanisms in mouse models: practical guidelines for systematically evaluating phenotypic changes in the diaphyses of long bones. J Bone Miner Res. 2015;30(6):951–66.

25. Turner CH, Burr DB. Basic biomechanical measurements of bone: a tutorial. Bone. 1993;14(4):595–608.

26. Guss JD, Horsfield MW, Fontenele FF, Sandoval TN, Luna M, Apoorva F, et al. Alterations to the Gut Microbiome Impair Bone Strength and Tissue Material Properties. J Bone Miner Res. 2017;32(6):1343–53.

27. Gilet V, Mabilleau G, Loumaigne M, Coic L, Vitale R, Oberlin T, et al. Superpixels meet essential spectra for fast Raman hyperspectral microimaging. Opt Express. 2024;32(1):932–48.

28. Li Z, Hardij J, Evers SS, Hutch CR, Choi SM, Shao Y, et al. G-CSF partially mediates effects of sleeve gastrectomy on the bone marrow niche. J Clin Invest. 2019;129(6):2404–16.

29. Pluskiewicz W, Buzga M, Holeczy P, Bortlik L, Smajstrla V, Adamczyk P. Bone mineral changes in spine and proximal femur in individual obese women after laparoscopic sleeve gastrectomy: a short-term study. Obes Surg. 2012;22(7):1068–76.

30. Scibora LM. Skeletal effects of bariatric surgery: examining bone loss, potential mechanisms and clinical relevance. Diabetes Obes Metab. 2014;16(12):1204–13.

31. Stein EM, Silverberg SJ. Bone loss after bariatric surgery: causes, consequences, and management. Lancet Diabetes Endocrinol. 2014;2(2):165–74.

32. Athonvarangkul D, Wysolmerski JJ. Crosstalk within a brain-breast-bone axis regulates mineral and skeletal metabolism during lactation. Front Physiol. 2023;14:1121579.

33. Miller SC, Bowman BM. Rapid inactivation and apoptosis of osteoclasts in the maternal skeleton during the bone remodeling reversal at the end of lactation. Anat Rec (Hoboken). 2007;290(1):65–73.

34. Garner SC, Peng TC, Hirsch PF, Boass A, Toverud SU. Increase in serum parathyroid hormone concentration in the lactating rat: effects of dietary calcium and lactational intensity. J Bone Miner Res. 1987;2(4):347–52.

35. Hernandez LL, Gregerson KA, Horseman ND. Mammary gland serotonin regulates parathyroid hormone-related protein and other bone-related signals. Am J Physiol Endocrinol Metab. 2012;302(8):E1009–15.

36. Lloyd JT, Alley DE, Hawkes WG, Hochberg MC, Waldstein SR, Orwig DL. Body mass index is positively associated with bone mineral density in US older adults. Arch Osteoporos. 2014;9:175.

37. Compston J. Obesity and fractures. Joint Bone Spine. 2013;80(1):8–10.

38. Inzana JA, Kung M, Shu L, Hamada D, Xing LP, Zuscik MJ, et al. Immature mice are more susceptible to the detrimental effects of high fat diet on cancellous bone in the distal femur. Bone. 2013;57(1):174–83.

39. Shen CL, Chen L, Wang S, Chyu MC. Effects of dietary fat levels and feeding durations on musculoskeletal health in female rats. Food Funct. 2014;5(3):598–604.

40. Bozkurt E, Kaya C, Omeroglu S, Guven O, Mihmanli M. The rapid effects of sleeve gastrectomy on glucose homeostasis and resolution of diabetes mellitus. Endocrinol Diabetes Metab. 2021;4(2):e00182.

41. Brinckerhoff TZ, Bondada S, Lewis CE, French SW, DeUgarte DA. Metabolic effects of sleeve gastrectomy in female rat model of diet-induced obesity. Surg Obes Relat Dis. 2013;9(1):108–12.

42. Cheah S, Gao Y, Mo S, Rigas G, Fisher O, Chan DL, et al. Fertility, pregnancy and post partum management after bariatric surgery: a narrative review. Med J Aust. 2022;216(2):96–102.

43. Gudbrandsen OA, Dankel SN, Skumsnes L, Flolo TN, Folkestad OH, Nielsen HJ, et al. Short-term effects of Vertical sleeve gastrectomy and Roux-en-Y gastric bypass on glucose homeostasis. Sci Rep. 2019;9(1):14817.

44. Harris DA, Mina A, Cabarkapa D, Heshmati K, Subramaniam R, Banks AS, et al. Sleeve gastrectomy enhances glucose utilization and remodels adipose tissue independent of weight loss. Am J Physiol Endocrinol Metab. 2020;318(5):E678–E88.

45. Micic DD, Toplak H, Micic DD, Polovina SP. Reproductive outcomes after bariatric surgery in women. Wien Klin Wochenschr. 2022;134(1-2):56–62.

46. Donnelly E, Meredith DS, Nguyen JT, Gladnick BP, Rebolledo BJ, Shaffer AD, et al. Reduced cortical bone compositional heterogeneity with bisphosphonate treatment in postmenopausal women with intertrochanteric and subtrochanteric fractures. J Bone Miner Res. 2012;27(3):672–8.

47. Farlay D, Rizzo S, Ste-Marie LG, Michou L, Morin SN, Qiu S, et al. Duration-Dependent Increase of Human Bone Matrix Mineralization in Long-Term Bisphosphonate Users with Atypical Femur Fracture. J Bone Miner Res. 2021;36(6):1031–41.

48. Forte YS, Renovato-Martins M, Barja-Fidalgo C. Cellular and Molecular Mechanisms Associating Obesity to Bone Loss. Cells. 2023;12(4).

49. Gagnon C, Schafer AL. Bone Health After Bariatric Surgery. JBMR Plus. 2018;2(3):121–33.

