## Supplementary figures and images for "Effects of vertical sleeve gastrectomy prior to pregnancy on bone mass, microarchitecture and material properties in the female rat"

### Supplemental Figure 1

#### Supplemental material

**Supplementary figure 1:** Schematic of the study design

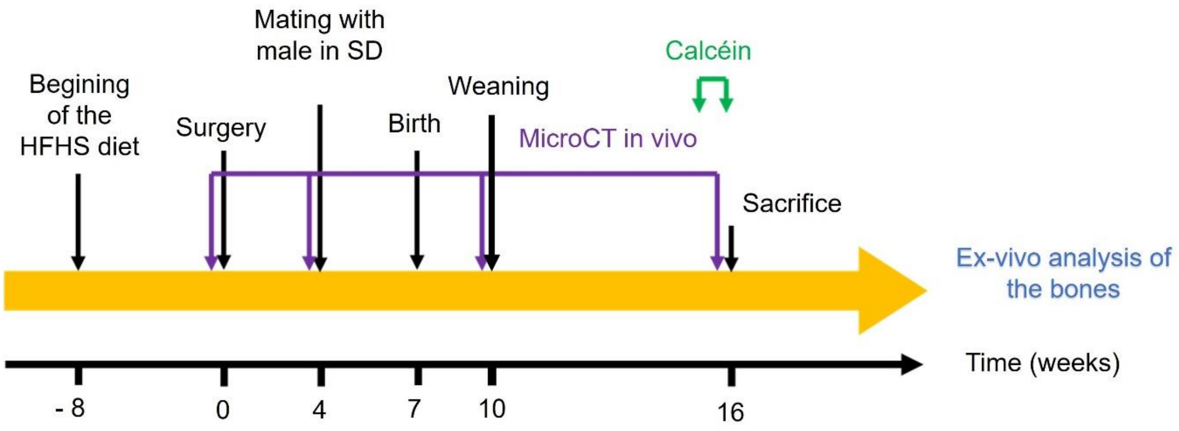
